# An experimental test of the relationship between melanism and desiccation survival in insects

**DOI:** 10.1101/012369

**Authors:** Subhash Rajpurohit, Lisa Marie Peterson, Andrew Orr, Anthony J. Marlon, Allen G Gibbs

## Abstract

We used experimental evolution to test the ‘*melanism-desiccation*’ hypothesis, which proposes that dark cuticle in several *Drosophila* species is an adaptation for increased desiccation tolerance. We selected for dark and light body pigmentation in replicated populations of *D. melanogaster* and assayed traits related to water balance. We also scored pigmentation and desiccation tolerance in populations selected for desiccation survival. Populations in both selection regimes showed large differences in the traits directly under selection. However, after over 40 generations of pigmentation selection, dark-selected populations were not more desiccation-tolerant than light-selected and control populations, nor did we find significant changes in carbohydrate amounts that could affect desiccation resistance. Body pigmentation of desiccation-selected populations did not differ from control populations after over 140 generations of selection. Our results do not support an important role for melanization in *Drosophila* water balance.

## INTRODUCTION

Pigmentation in insects is extremely diverse, both among and within species (Mani 1968; Majerus 1998; True 2003; Gray and McKinnon 2007; Kronforst et al. 2012). In addition to differences in color, insects can differ in the deposition of melanin, a dark polymer of dopa derivatives. Several adaptive hypotheses have been proposed for variation in melanization (True 2003; Wittkopp and Beldade 2009; Kalra et al. 2014). These include behavioral benefits (crypsis, sexual selection, etc.) and physiological benefits, including thermoregulatory capacity and resistance to abrasion, ultraviolet radiation, infection and desiccation (Johnson et al. 2011; Bastide et al. 2014; Rajpurohit et al. 2008a). In recent years, several research groups have used *Drosophila* to investigate the functional significance of melanization, particularly in the context of desiccation stress and water balance. Water balance is a general physiological problem for insects, because their large surface area:volume ratio makes insects susceptible to water loss through the cuticle. Melanization may reduce water loss by making the cuticle thicker or more hydrophobic (Gibbs and Rajpurohit 2010).

The genus *Drosophila* provides an excellent system in which to examine the function of melanization. Species differ widely in their body pigmentation, and pigmentation mutants have been identified in several species. For example, *ebony* mutants of *D. melanogaster* are more resistant to desiccation stress than wildtype flies, whereas *yellow* mutants of multiple species are less desiccation resistant (Kalmus 1941). Within species, darker populations tend to be more desiccation resistant (Brisson et al. 2005; Rajpurohit et al. 2008b), and, within populations, darker pigmentation is also associated with increased desiccation resistance (Parkash et al. 2009).

A series of recent studies have examined parallel clines in desiccation tolerance and body melanisation in several *Drosophila* species from the Indian subcontinent (see Parkash 2010; Rajpurohit et al. 2013). Populations of *D. melanogaster* from higher (and dryer) latitudes and altitudes on the Indian subcontinent are darker and more resistant to desiccation (Parkash et al. 2008, Rajpurohit et al. 2008a; Rajpurohit et al. 2013; Rajpurohit and Nedved 2013), suggesting that differences in pigmentation are functionally significant. This idea is supported by the findings that darker phenotypes of *D. melanogaster* and other species lose water less rapidly than lighter phenotypes (Parkash 2010; Ramniwas et al. 2013). A potential mechanistic explanation for these correlations is the hydrophobic nature of melanin. Like epicuticular hydrocarbons, melanin may decrease the permeability of the cuticle to water.

A central problem for studies of clinal variation is that environmental factors often co-vary, making it difficult to distinguish which factor (or combination thereof) is responsible for the cline. For example, a recent synthesis of clinal variation in Indian populations of *Drosophila* (Rajpurohit et al. 2013) found that desiccation resistance was most highly associated with the coefficient of variation in monthly temperature, which was highly correlated with mean annual temperature, mean relative humidity, and the coefficient of variation of monthly relative humidity. Thus, parallel clines in traits may arise from independent selection exerted by parallel clines in environmental variables. For example, selection for reduced water loss in dry environments may coincide with selection for thermoregulatory capability, resulting in co-evolved differences in these traits.

Experimental evolution provides a means to manipulate environmental factors independently and to rigorously test whether correlated variation observed in natural populations has a physiological or genetic basis. Direct comparison of laboratory and natural systems provides the opportunity to identify and test hypotheses regarding natural selection in the field (Gibbs 1999; Gibbs and Gefen 2009; Dykhuizen and Dean 2009; Rose et al. 1996; Folk and Bradley 2005). If laboratory and comparative studies provide similar results, this is corroborative evidence that selection is acting as we thought in nature (Lynch 1992). When different results are obtained, then something may be missing in our understanding of one or both environments (Gibbs 1999).

In this study, we used experimental evolution to test the hypothesis that melanism and desiccation tolerance are functionally associated in *D. melanogaster*. Previous studies have demonstrated that natural populations of *D. melanogaster* harbor significant genetic variation for both traits. Pigmentation and desiccation tolerance each respond rapidly to selection in the laboratory (pigmentation: Rajpurohit and Gibbs 2012; desiccation tolerance: Rose et al. 1992; Archer et al. 2006). We reasoned that selecting populations for darker or lighter pigmentation should result in populations with greater or lesser desiccation tolerance, respectively. Conversely, selection for increased desiccation tolerance should result in darker populations of *Drosophila*. Our results contradict these predictions. Desiccation-selected flies were not darker than controls, and pigmentation-selected populations exhibited relatively small differences in desiccation resistance that were not consistent with our predictions. We discuss potential correlated responses to selection on other traits associated with water balance, such as body size and carbohydrate content.

## MATERIAL AND METHODS

### Selection for body melanization and desiccation tolerance

All populations of *D. melanogaster* were maintained at 24°C under constant light. Desiccation-selected lines were founded from ~400 females collected in Terhune Orchard, New Jersey, USA in 1999, and pigmentation-selected lines were founded from a population (~400 individuals) collected from Las Vegas, Nevada, USA in 2008. The population selection and maintenance procedures have been described elsewhere (Gefen et al. 2006; Rajpurohit and Gibbs 2012).

Selecting for pigmentation entailed artificial selection in the laboratory (Rose *et al.* 1996), in which the darkest or lightest 10% females, as chosen by the primary author, were allowed to reproduce each generation. 200 one-week-old females were randomly selected from each population, and the 20 darkest or lightest were allowed to lay eggs for the next generation. Three replicate dark-selected (D_PIG_) and light-selected (L_PIG_) populations were created from the initial founding population, along with three control (C_PIG_) populations, for which 20 breeding females were selected randomly from the population each generation (Rajpurohit and Gibbs 2012).

To select for desiccation resistance, we used natural selection in the laboratory (Rose et al. 1996). Three replicated populations (D) were selected for desiccation tolerance, and three control populations were maintained without desiccation stress (F, continuous access to food and water). For desiccation selection, ~10,000 flies were exposed to low humidity conditions (no food, in the presence of desiccant) each generation, and the 1000-1500 individuals that survived the longest were allowed to recover and produce offspring. This procedure was followed for 30 generations; subsequent generations of D populations were subjected to desiccation stress for 24 hours each generation, a condition causing 100% mortality in F populations (Gefen et al. 2006; confirmed in experiments described below). Desiccation-selected populations had undergone ~140 generations of selection (30 generations of strong selection, ~110 generations of relaxed selection) when experiments were performed.

### Egg collection for experimental assays

Adults from each of the populations were transferred to empty 175-ml bottles for one hour. The bottles were covered with a 35x10 mm Petri dish containing grape agar as a substrate for egg laying. Sets of 60 eggs were collected in replicates and placed in food vials containing approximately 10 ml of cornmeal-yeast sucrose media. The vials were incubated at 24°C with constant light. To avoid potential parental effects, the populations were kept off selection for one generation before performing any analyses.

### Desiccation resistance

Four-to-five day old flies were transferred to empty vials individually, and restricted to the lower half of the vials by a foam stopper. Silica gel was then added above the stopper to maintain low humidity, and the vial was sealed with Parafilm. Mortality was recorded at hourly intervals until all flies were dead.

### Body tergite pigmentation scoring and area measurements

For the measurements of body tergite pigmentation and size, we used whole mount abdomens which were prepared on transparent glass slides. The mounted abdomens were imaged using a Nikon digital camera attached to a dissecting microscope. The images were analyzed using ImageJ software (http://rsbweb.nih.gov/ij/). We collected data for abdominal pigmentation and total dorsal abdominal area based on the method developed by Rajpurohit and Marlon (2011). The measurements were calibrated using a glass marked scale. A scale image was taken before the sample images taken (without changing magnification between the scale image and sample images). While collecting data a scale unit was set, and images were processed with that base scale. These measurements used seven-to-ten day old flies.

### Wet mass, dry mass and water content

Four-day old flies were weighed on a Cahn C-30 microbalance. To estimate wet weight, the flies were frozen at -20 °C and weighed immediately after removal from the freezer. Dry weights were measured as the weight after drying at 50 °C overnight. Total body water content was estimated as the difference between masses before and after drying at 50 °C.

### Carbohydrate assays

Newly eclosed flies, from all 9 experimental groups of pigmentation selection types, were collected as described above and immediately frozen at −20 °C. After thawing, the flies were sexed, homogenized in 200 μl 0.05% Tween-20, and incubated at 70°C for 5·min. The samples were then centrifuged for 1·min at 16·000·***g***, and the supernatants removed and frozen until measurements. For carbohydrate measurements, we followed the methods used by Gefen et al. (2006).

### Flow-through respirometry

Water-loss rates and metabolic rates were measured using flow-through respirometry (TR-2 respirometer; Sable Systems, Las Vegas, Nevada, USA). Groups of 10-20 flies were placed in 5ml glass/aluminum chambers, and dry CO_2_-free air was pumped through the chambers at a flow rate of 50 ml min^-1^ to an LI-6262 infrared CO_2_ sensor (Li-Cor Biosciences, Lincoln, Nebraska, USA). Metabolic and water-loss rates were calculated from CO_2_ and water vapor released by flies into the air stream. The humidity sensor was calibrated by injection of small drops of water (0.5–3.0 nl) into the air stream, and the CO_2_ detector was calibrated according to the manufacturer’s instructions using 100 ppm span gas. Datacan V software (Sable Systems, NV, USA) was used for data collection and analysis.

### Statistical analyses

We used Statistica v7.1 to analyze our data. Desiccation- and pigmentation-selected populations (and their respective controls) were analyzed separately. We used mixed-model analyses of variance (ANOVA), with selection and sex as fixed main effects and replicate population as a random variable nested within selection treatment. Desiccation-resistance data were analyzed using a log-ranks test, with censoring for missing data points (mostly from the end of the experiment).

## RESULTS

### Pigmentation

Body tergite melanization for pigmentation- and desiccation-selected populations are shown in Supplementary Fig. 1 and Fig. 1, respectively. Populations selected for body pigmentation (D_PIG_, C_PIG_ and L_PIG_) showed a significant response to selection (F_2,161_=99.91; P<0.00002; see Table S1). Although females were the target of selection in pigmentation-selection regime, males also evolved differences in pigmentation (only female phenotypes are shown in Supplementary Fig. 1). Besides selection and sex as significant main effects, selection*sex interaction effects were also significant (F_6,161_=3.24; P<0.004; Table S1). No differences in abdominal tergite pigmentation were detected between desiccation-selected and control populations after 140 generations of selection (Table 1; F_1,198_=1; P<0.37).

**Figure 1:**
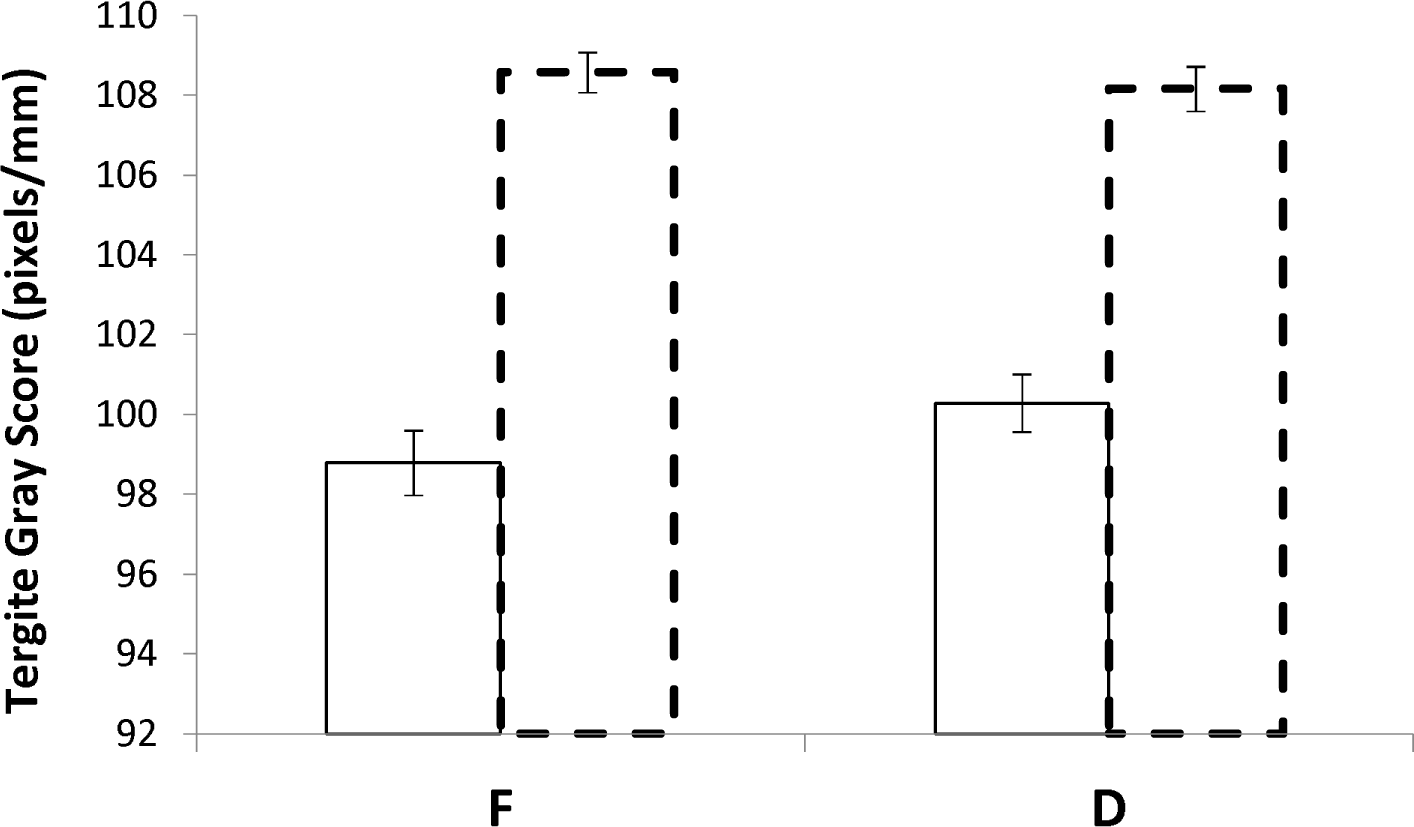
Pigmentation status (as gray scale score) in D and F populations (>140 generations of artificial selection). Solid line bars and dotted line bars represent male and female data respectively.

**Table 1:** Nested ANOVA results for pigmentation status in D and F populations.

| Parameters | SS | d.f. | MS | F | *p* |
| --- | --- | --- | --- | --- | --- |
| Selection | 17 | 1 | 17 | 1 | 0.36366 |
| Replicate(Selection) | 64 | 4 | 16 | 1.4 | 0.366663 |
| Sex | 4067 | 1 | 4067 | 364 | **0.000043** |
| Replicate(Selection*Sex) | 45 | 4 | 11 | 0.5 | 0.752936 |
| Selection*Sex | 37 | 1 | 37 | 3.3 | 0.142661 |
| Error | 4636 | 198 | 23 |  |  |

### Desiccation resistance

Differences observed in desiccation resistance for L_PIG_, C_PIG_ and D_PIG_ populations are shown in Fig. 2. An ANOVA revealed significant effects of replicate (selection) (Table 2; F_6,200_=78, P<0.00002). As expected, significant differences in desiccation tolerance between the sexes were found in pigmentation-selected populations. Males died earlier than females under desiccating conditions (Fig. 2).

**Figure 2:**
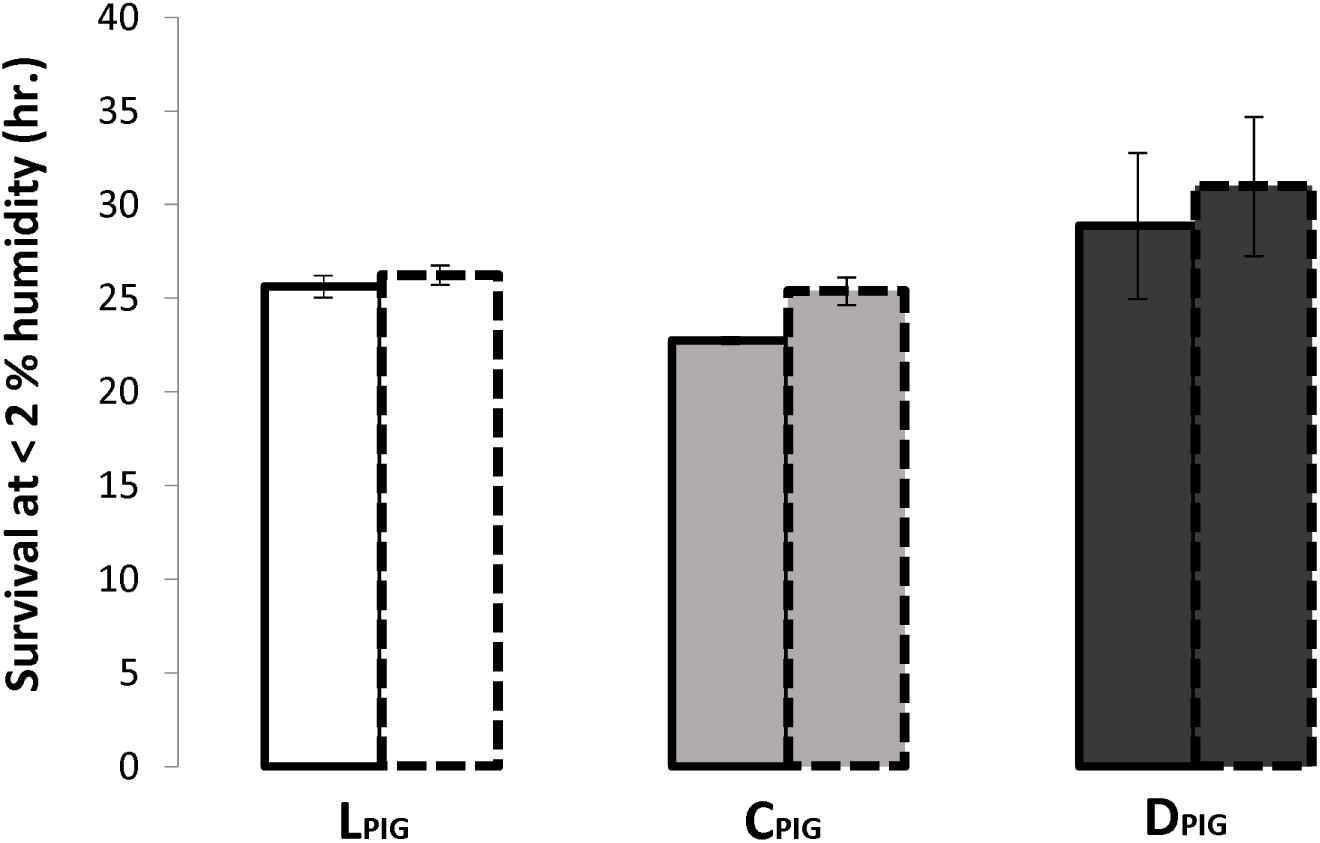
Survival hours under desiccating conditions (without food and water; <2% humidity conditions) in L_PIG_, C_PIG_ and D_PIG_ populations. Solid line bars and dotted line bars represent male and female data respectively.

**Table 2:** Nested ANOVA results for desiccation tolerance in pigmentation-selected populations and controls.

| Parameters | SS | d.f. | MS | F | *p* |
| --- | --- | --- | --- | --- | --- |
| Selection | 1277.4 | 2 | 638.7 | 1.8143 | 0.241968 |
| Replicate(Selection) | 2112.5 | 6 | 352.1 | 77.963 | **0.00002** |
| Sex | 172.7 | 1 | 172.7 | 38.2267 | **0.000821** |
| Replicate(Selection*Sex) | 27.1 | 6 | 4.5 | 0.4993 | 0.808452 |
| Selection*Sex | 41 | 2 | 20.5 | 4.5343 | 0.063098 |
| Error | 1809 | 200 | 9 |  |  |

In assays of desiccation-selected (D) populations, most fed control (F) flies had died within 15 hours under desiccating conditions, when the earliest mortality occurred for D flies. Some survival data were missing at the tails of the F and D survival curves, so we compared each replicate D and F population to all of the other populations using log-ranks tests, with a sequential Bonferroni correction for multiple comparisons (Table S2). All D replicates survived longer than all F replicates. Every pair-wise comparison of a D and an F population was statistically significant, while no comparisons between replicates within a selection treatment were significant (Table S2). When replicate populations were pooled, desiccation resistance of D flies was significantly greater than that of F flies (P < 10^-5^; not shown).

### Wet and dry mass, carbohydrate and total water content

Pigmentation-selected populations did not differ in carbohydrate content from their controls or from each other (Fig. 3; Table 3). The direction of carbohydrate content was in the following order: L_PIG_>D_PIG_>C_PIG_. Nested ANOVA revealed significant differences in total carbohydrate content across replicate types when they were nested within selection*sex (Table 3; F_6,124_=3.51; P<0.003). Females accumulated more carbohydrate than males (Fig. 3, Table 3).

**Figure 3:**
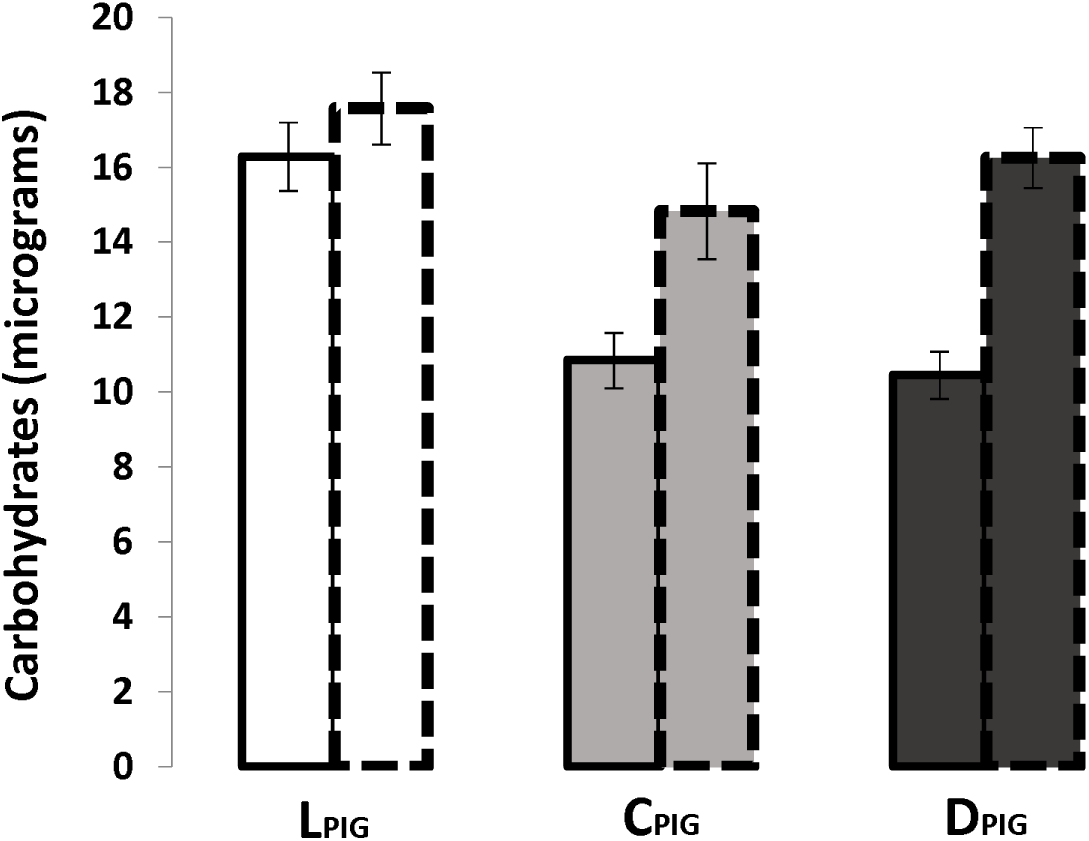
Status of total carbohydrates in L_PIG_, C_PIG_ and D_PIG_ populations. Solid line bars and dotted line bars represent male and female data respectively.

**Table 3:** Nested ANOVA results for carbohydrate status in pigmentation-selected populations and controls.

| Parameters | SS | d.f. | MS | F | *p* |
| --- | --- | --- | --- | --- | --- |
| Selection | 491.18 | 2 | 245.59 | 2.2273 | 0.189008 |
| Replicate (selection) | 662.28 | 6 | 110.38 | 2.218 | 0.177593 |
| Sex | 457.45 | 1 | 457.45 | 9.2022 | **0.022969** |
| Replicate (selection*sex) | 298.59 | 6 | 49.77 | 3.5184 | **0.003002** |
| Selection*Sex | 135.36 | 2 | 67.68 | 1.3611 | 0.32547 |
| Error | 1753.89 | 124 | 14.14 |  |  |

We also analyzed wet mass, dry mass and total water content at the time of eclosion in pigmentation-selected populations. Nested ANOVA revealed no difference in wet mass in D_PIG_, C_PIG_ and L_PIG_ populations (selection F_2,520_=1.759; P<0.25; Table S3A), but a significant difference was observed in dry mass (selection F_2,520_=7.18; P<0.03; Table S3B). Male to female differences in wet mass, dry mass and total water content were significant (data not shown). The D populations had higher amounts than F populations of carbohydrates and total water content in both males and females (see Gefen et al. 2006).

### Metabolic rate and water loss rate

We found no differences in metabolic rate and water-loss rate among D_PIG_, C_PIG_ and L_PIG_ flies (MR F_2,74_=0.5; P>0.6 and WLR F_2,74_=5; P>0.6; Table S4), although significant differences appeared among replicates (nested within selection). In contrast, desiccation-selected D flies lost water significantly slower than fed F controls (F_1,60_=5.2, P<0.03; Fig. 4; Table S5).

**Figure 4:**
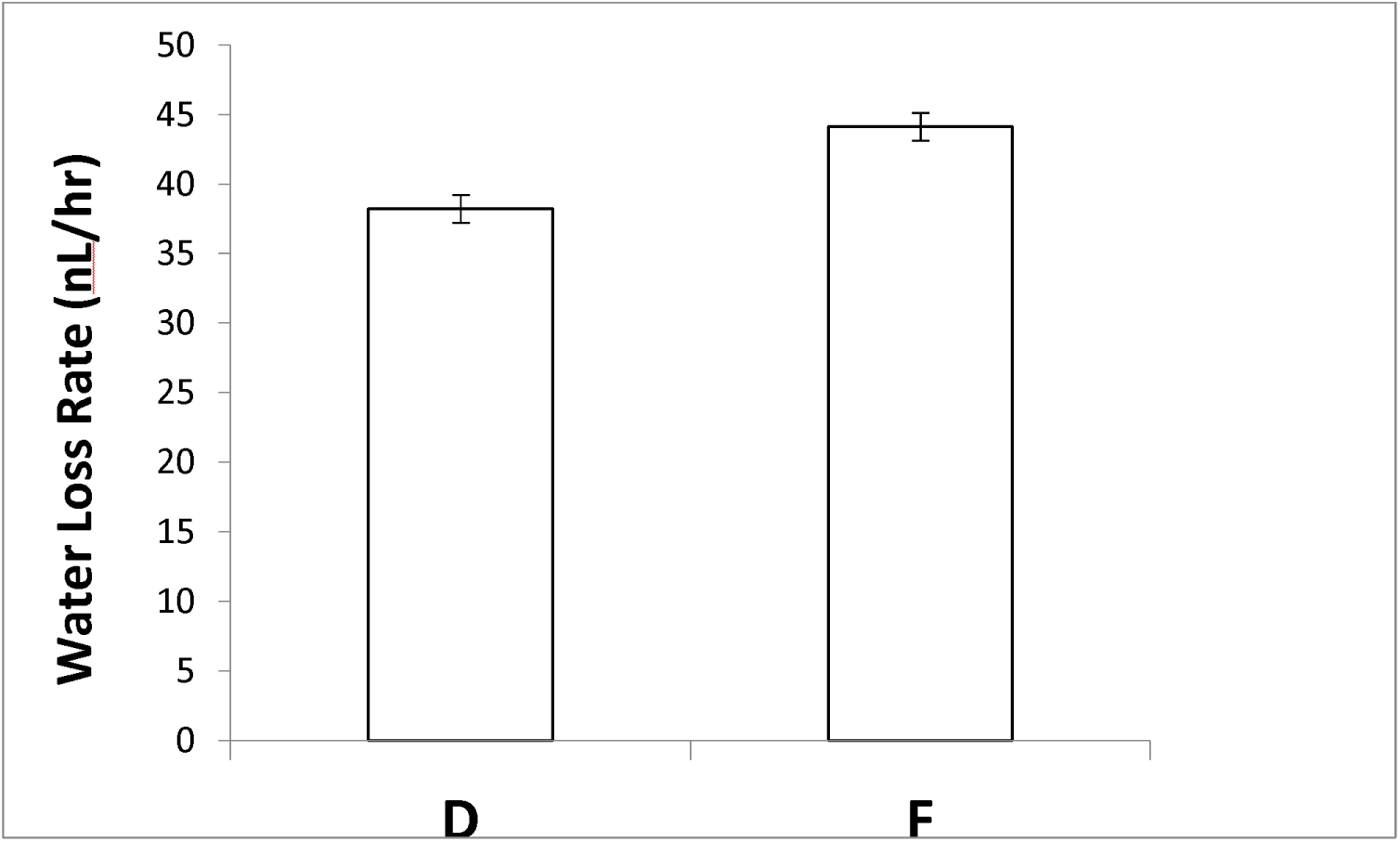
Water loss rate in D and F populations. Only female data is shown here.

## DISCUSSION

Abdominal pigmentation and trident (a thoracic pigmentation pattern) show similar latitudinal and altitudinal clines in several *Drosophila* species (David et al. 1985; Munjal et al. 1997). On the Indian subcontinent, darker populations with a more pronounced trident are observed at higher latitudes and elevations (Munjal et al. 1997). Darker abdominal and thoracic (trident) pigmentation will both contribute to an overall darker body. This may increase absorption of solar radiation, so that darker flies can remain active at lower environmental temperatures (Gibert et al. 1998; Parkash et al. 2010; True 2003). In several insects, a significant effect of pigmentation on thermoregulation has been reported (Watt 1969; Brakefield and Willmer 1985). However, the importance of pigmentation for temperature regulation in drosophilids and other small insects remains to be established experimentally. Willmer and Unwin (1981) reported that insects the size of *Drosophila* do not attain temperatures even 1°C above ambient, whereas large (>100 mg), dark insects with poorly reflective bodies can achieve body temperatures >10°C above ambient (Prange 1986). Hirai and Kimura (1997) found that black morphs of *D. elegans*, which is larger than *D. melanogaster*, were slightly (0.26°C) warmer than brown morphs under irradiation. Thus, the thermal significance of melanization is size-dependent, and *D. melanogaster* is too small for melanization to affect its body temperature.

Pigmentation may have adaptive significance related to other physiological processes, such as desiccation resistance, protection against ultraviolet radiation (True 2003) or resisting infection (Dombeck and Jaenike 2004). At the population level, darker pigmentation is correlated with reduced water loss and increased desiccation resistance in several *Drosophila* species (Brisson et al. 2005; Rajpurohit et al. 2008a; Kalra et al. 2014). Within populations, darker pigmentation is also associated with increased desiccation resistance (Parkash et al. 2009). Laboratory studies in *D. melanogaster* and *D. polymorpha* show differences in desiccation resistance between color morphs (Kalmus 1941; Brisson et al. 2005). In contrast, natural populations of *D. americana* from a longitudinal cline in North America, are darker in more humid areas (Wittkopp et al. 2011). This suggests that selection promoting the pigmentation cline in *D. americana* might be different from that in other *Drosophila* species. In general, however, across many *Drosophila* species and continents, melanism and desiccation resistance are correlated. Melanin’s hydrophobic nature is consistent with a reduction in cuticular permeability, making this an attractive physiological hypothesis.

If melanism and desiccation resistance are indeed mechanistically linked through differences in cuticular permeability, then selection on either trait should result in correlated responses in the other. We performed two complementary selection experiments, which together suggest that the *melanism-desiccation* hypothesis is incorrect (see Fig. 1 & Table S4). However, survival under desiccating conditions is a function of multiple physiological characters. Our studies reveal that some other characters have undergone correlated responses to selection in both pigmentation- and desiccation-selected populations.

Our strongest evidence that melanism has little effect on cuticular water loss is provided by the pigmentation-selected populations. Despite clearly visible differences in appearance, light- and dark-selected flies did not differ in overall water-loss rates (Table S4). Cuticular transpiration and respiration are the primary routes for water loss from insects (Chown et al. 2006; Quinlan and Gibbs 2006). Despite relative inactivity, however, desiccation-selected flies do not have lower respiratory water-loss rates than controls (Williams and Bradley 1998). Metabolic rates did not differ among pigmentation-selected populations and their controls (Table S4), suggesting that differences in respiratory water loss did not affect overall water-loss rates.

Epicuticular hydrocarbons (HC) provide an important barrier to cuticular transpiration in insects (Gibbs and Rajpurohit 2010), and it is possible that selected populations differed in the amount and/or composition of HC (Foley and Telonis-Scott 2011). Such differences have been implicated in latitudinal clines in Indian populations of other drosophilids (Kalra et al. 2014). However, longer-term (160 generations), more stringent desiccation selection than performed in this study yielded only minor HC differences (Gibbs et al. 1997). Inter-specific and acclimatory studies reveal no consistent relationships between water-loss rates and HC (Gibbs et al. 1998, 2003; Gibbs and Matzkin 2001). Although we cannot rigorously exclude HC differences between selected populations and controls in this study, previous research suggests that these are not important factors in desiccation resistance of either set of selected and control populations.

In addition to water loss, the ability of animals to survive desiccating conditions depends on how much water is contained within the animal when desiccation begins, and how well the animal tolerates low water content (Gibbs and Gefen 2009). In *Drosophila*, there is little support for the latter mechanism in either desiccation-selected or natural populations (Gibbs et al. 1997; Gibbs and Matzkin 2001).

Water content includes both bulk (pre-formed) water and water available from metabolism of lipids, glycogen or proteins. In the case of lipids, these are distinct pools, as lipid droplets in cells bind very little water. The situation is more complicated for glycogen and protein, each of which can bind several times their mass in water of hydration. When these macromolecules are metabolized, their bound water is released and is available to replace bulk water lost from the animal. Desiccation-selected populations of *D. melanogaster* contain more glycogen than control populations (Gibbs et al. 1997; Chippindale et al. 1998; Folk et al. 2001; Gefen et al. 2006). *Drosophila* species in general tend to metabolize glycogen under desiccating conditions (Marron et al. 2003; Parkash et al. 2012), consistent with the release of bound water when it is required. Glycogen storage is not incompatible with storing extra bulk water, as indicated by the 3-fold greater hemolymph volumes of desiccation-selected flies relative to controls (Folk et al. 2001).

Previous work has shown that D populations contain more carbohydrate than their F controls (Gefen et al. 2006). Flies reared on poor quality food as larvae are lighter in color than well-fed controls (Shakhmanstsir et al. 2014). This suggests a tradeoff between allocation to pigmentation and other organismal requirements that is revealed under low nutrition conditions. This led us to investigate whether pigmentation selection resulted in a correlated response in carbohydrate storage, with higher storage in L_PIG_ flies potentially counteracting higher water-loss rates. However, carbohydrate levels did not follow a consistent pattern with melanism (Fig. 3; Table 3), leading us to reject this idea.

### Conclusions

We performed complementary selection experiments in *Drosophila* to test the hypothesis that melanism and desiccation resistance are functionally linked. Neither experiment yielded results consistent with our predictions, so we reject this simple hypothesis. We note, however, that other physiological and behavioral variables affect water loss, and that melanism has multiple other potential functions in *Drosophila*. Tradeoffs between these functions and pleiotropic effects of genes involved in melanin metabolism are likely to result in complex interactions.

## ACKNOWLEDGEMENTS

This work was supported through National Science Foundation grant 0723930.

---

**Supplementary Figure 1:**
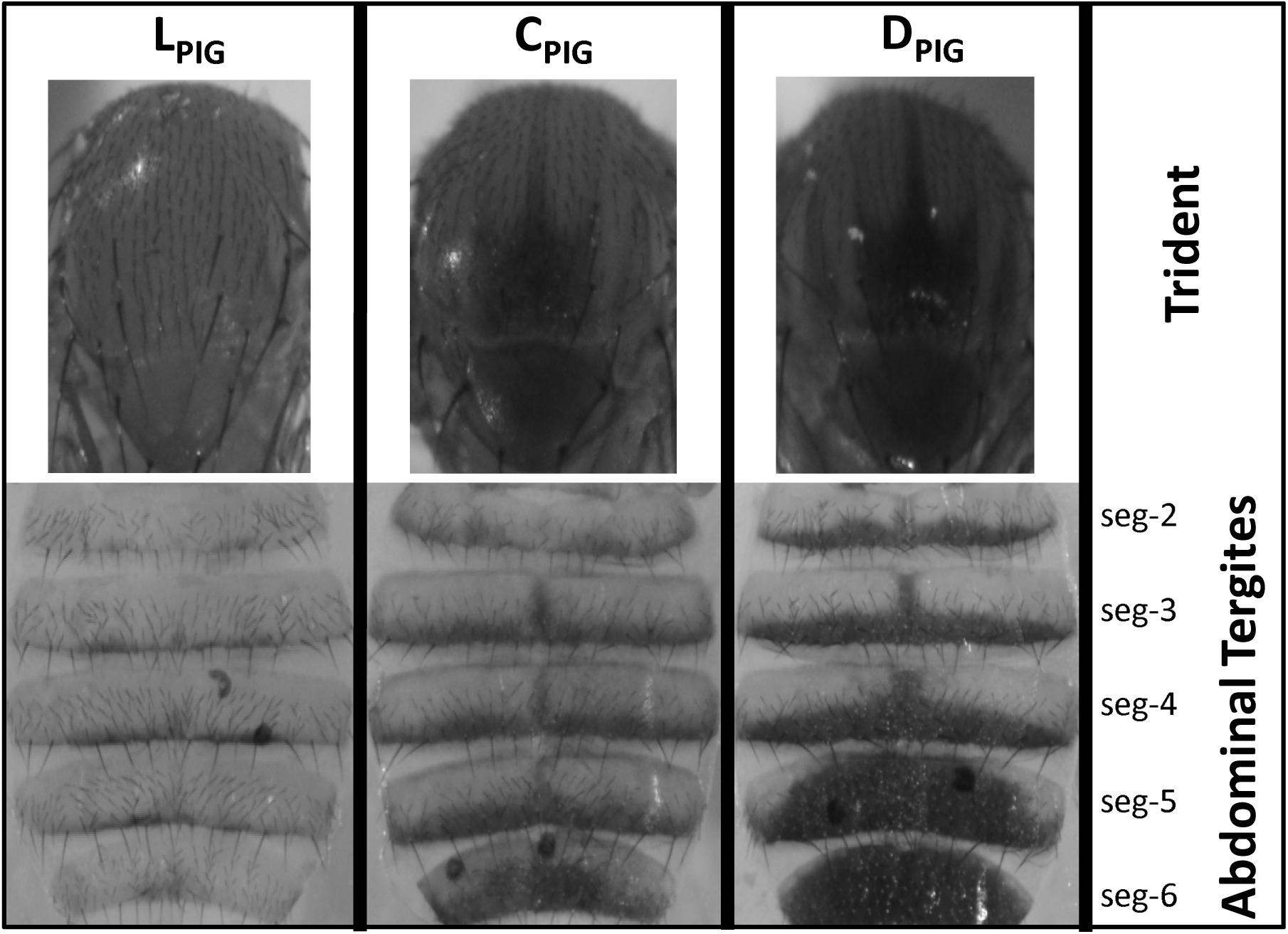
Trident and abdominal tergite pigmentation status (>40 generations of artificial selection) in L_PIG_, C_PIG_ and D_PIG_ populations. Only female images are shown here. Males also showed parallel response. Please neglect two black dotted structures under abdominal tergite. These are the parts of female reproductive tract while flattening dorsal segments.

